# Preparation of H-Ras GTPase conjugated to lipid nanodiscs for NMR spectroscopy

**DOI:** 10.1101/181032

**Authors:** Elizaveta A Kovrigina, Sneha Shah, Evgenii L Kovrigin

## Abstract

Ras GTPase is a peripheral membrane protein central to cellular signaling of growth and proliferation. Membrane attachment is critical for a range of Ras activities, therefore, ability to make faithful *in-vitro* samples of a mem-brane-bound Ras for detailed biophysical studies is a highly desirable goal. In this manuscript, we are describing preparation of a large-scale sample of isotopically labeled H-Ras conjugated to lipid nanodiscs. We demonstrate that the Ras-nanodisc sample is fairly stable to allow for a range of Nuclear Magnetic Resonance (NMR) and other biophysical measurements. The need to achieve a homogeneous protein-nanodisc ratio is also emphasized.

## Introduction

Ras is a small monomeric GTPase acting as a molecular switch in a range of critical cellular processes^*1*^. The protein consists of a well conserved G domain (homologous to the *α* subunit of the trimeric G proteins) and a lipidated C-terminal peptide, which is highly variable among the three isoforms of Ras, H, N, and K. The switch-like behavior is a function of a the G domain, which adopts the distinct conformations in the GTP-bound (“on”, active) or GDP-bound (“off”, inactive) states^*2*^. The C-terminal peptide (hyper-variable region, HVR) is anchored to the membrane with post-translationally attached lipid chains and is responsible for trafficking Ras to the specific plasma membane localizations^*3, 4*^. Binding partners of Ras (effectors) interact with the GTP-bound conformation of Ras through the so-called “effector interface” of the G domain^*2, 5*^. Most effectors require Ras attached to the membrane for full activity; many would bind to the Ras-GTP with truncated HVR in solution but not be activated (for a review see Henis, 2009^*6*^). Therefore, the comprehensive biophysical and structural studies of Ras interactions with its signaling binding partners must necessarily use Ras attached to the lipid bilayer.

Preparation of model bilayers correctly representing a flat surface of a cellular membrane yet amenable for structural biology studies has been one of the main bottlenecks in membrane protein research. Most of the published reports utilized detergent micelles or small lipid vesicles (SUV) to solubilize membrane-bound proteins (for example, see^*7-10*^). An important breakthrough occurred when the phospholipid bilayer nanodiscs have been introduced^*11, 12*^. These nanodiscs resemble the naturally occurring High-Density Lipoprotein particles (HDL) with a discoidal lipid bilayer surrounded by a matrix scaffold protein derived from apolipoprotein I (Apo-I). With the easily controlled size distribution and straightforward preparation it is possible to make nanodisks of various sizes from 12 nm diameter (ca. 240 kDa)^*12, 13*^ to 6 nm (ca. 54 kDa) ^*14*^ ideally suited for studies of membrane proteins. Recent advances included use of cirularized nanodiscs that result in extremely homogeneous nanodisc preparations^*15*^.

In this manuscript, we described preparation of the mem-brane-bound Ras sample conjugated to lipid nanodiscs and its preliminary characterization with the Nuclear Magnetic Resonance (NMR) spectroscopy.

## Preparation and characterization

Full-length lipidated Ras proteins may be expressed and isolated from mammalian or insect cells with native post-translational lipidation and carboxymethylation of the C-terminal peptide^*16*^. The high-yield preparations K-Ras (which carries a single farnesyl chain attachment) have been reported^*17, 18*^, but preparation of native H- and N-Ras proteins has been more difficult due to their multiple lipidation motifs. An alternative way to prepare the H and N-Ras proteins is through the semisynthetic approach: by joining a recombinant G domain and the synthetic lipidated peptide^*19*^. Because, by either methods, it is relatively difficult to produce large protein samples required for structural biology and biophysical studies, a simpler alternative was proposed: tethering Ras to the membrane surface through a non-natural lipid anchor^*20-22*^. In this model (Figure 1), the protein construct is stably associated with the bilayer through the irreversible conjugation of the C-terminal cysteine to the thiol-reactive derivative of phosphatidyl ethanolamine: 1,2-dioleoyl-sn-glycero-3-phosphoethanolamine-N-[4-(p-maleimidomethyl) cyclohexane-carboxamide] (PE-MCC).

**Figure 1.**
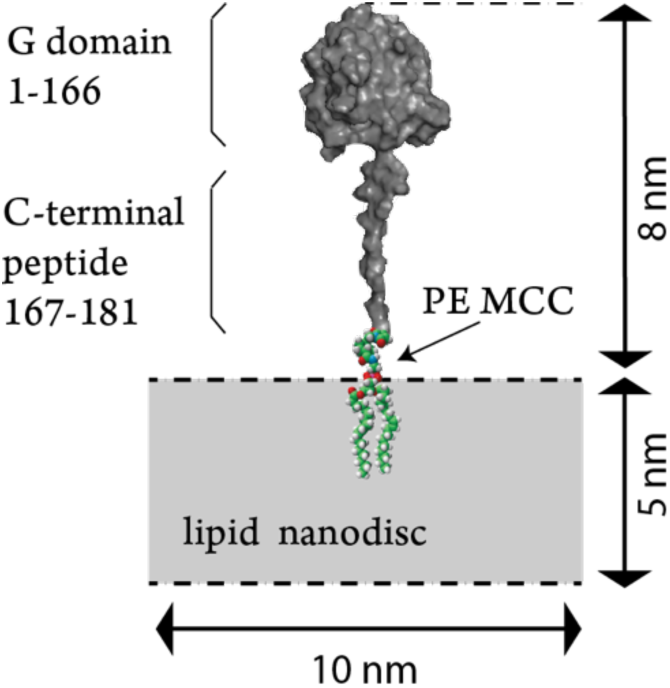
A model of the H-Ras 1-181C tethered to the PE MCC lipid. The model is approximately to scale. The gray rectangle depicts a cross-section of a lipid nanodisc. Ras is shown with the surface representation. G domain model is based on H-Ras 1-166 crystal structure PDBID 5P21. The C terminal peptide 167-181 was added in Pymol and modeled in the fully extended conformation.

To make the H-Ras sample conjugated to a lipid bilayer, we expressed and purified the construct that includes residues 1-181 of H-Ras ending at the fist cysteine residue (C181) of the hypervariable region. The overall expression and refolding procedure followed the published protocols^*23, 24*^. The Cys181 is the point of the first palmitoyl group attachment, one of the three lipids anchoring the native full-length H-Ras at the membrane^*25*^. Cysteine side chain irreversibly reacts with maleimide group on the PE MCC permanently attaching the PE phospholipid to the C-terminal of the Ras poly-peptide. The additional mutation, C118S, was introduced in H-Ras to remove the only exposed cysteine on the G domain, which may spuriously react with the maleimide group. The C118S mutation has been shown not to have any adverse effect on H-Ras function^*26*^. Figure 2 demonstrates the MALDI-TOF mass spectrum of the C118S H-Ras expression construct, residues 1-181 (“Ras” in the following text).

**Figure 2.**
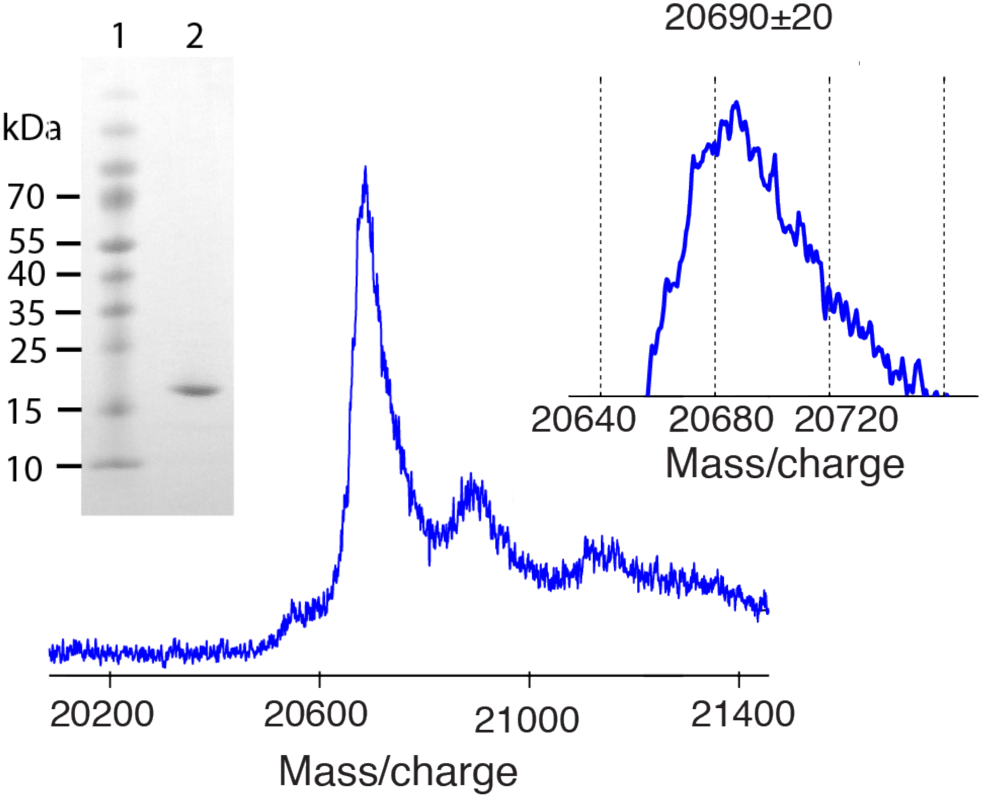
MALDI-TOF of the uniformly ^15^N-labeled C118S H-Ras, residues 1-181. (Left inset) SDS-PAGE of the purified protein: lane 1, markers; lane 2, protein sample prepared for conjugation with nanodiscs. (Middle) MALDI-TOF analysis of the sample from lane 2. Matrix: sinapinic acid. The tallest peak, H-Ras; minor peak to the right, sinapinic acid adduct. (Right inset) Enlarged view of the Ras peak maximum. Calculated molecular weight of ^15^N-uniformly labeled C118S H-Ras 1-181 is 20,678 Da.

To create 10-nm lipid nanodiscs for conjugation with Ras, we followed a standard protocol utilizing bacterially expressed matrix scaffold protein MSP1D1^*27*^ and the lipid mixture containing 80% DPPC, 10% DPPG and 10% PE MCC. The gel-filtration elution profile of the nanodisc sample (Figure 3, black trace) reveals a single major peak with the elution time corresponding to expected 164 kDa (two MSP1D1 molecules and ca. 160 lipids for the 10 nm nanodisc). Conjugation of Ras to maleimide-functionalized nanodiscs was performed by mixing equimolar quantities of the nanodiscs and Ras followed by incubation at the room temperature overnight (18 hours). At the end of incubation, the unreacted maleimide groups were quenched with beta-mercaptoethanol.

**Figure 3.**
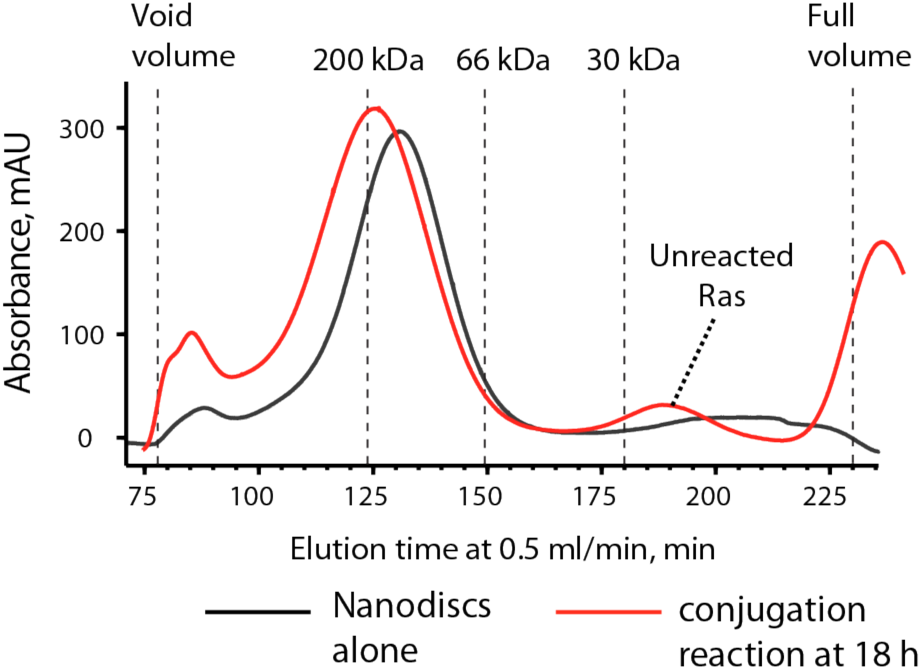
Gel-filtration profiles of a nanodisc sample (black) and the Ras-nanodisc conjugation reaction mixture after 18 hours of incubation at room temperature (red). Gel-filtration was performed on the XK16/70 column packed with Ultrogel Aca34 (Pall; 20-340 kDa separation range). Elution times of the molecular weight standards are shown by vertical dotted lines.

The gel-filtration profile of a reaction mixture is shown in Figure 3 as a red trace. The Ras-nanodisc conjugate elutes as the higher molecular weight species (ca. 200 kDa) with the residual unreacted Ras appearing in 20-30 kDa range. The calculated molecular weight of the Ras-nanodisc complex is approximately 184 kDa. Non-spherical shape of the assembly was expected to result in the somewhat greater hydrodynamic radius. The SDS PAGE analysis (Figure 4) confirms formation of the Ras-nanodisc conjugate clearly showing both MSP1D1 and Ras present in the 200 kDa peak fraction, lane 3.

**Figure 4.**
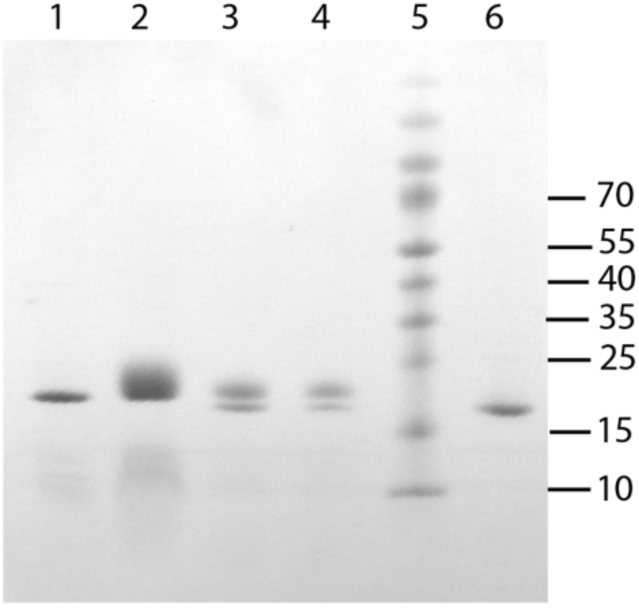
SDS-PAGE of proteins and nanodisc samples: lane 1, purified MSP1D1 used for nanodisc preparation; lane 2, purified sample of nanodiscs (from the 160 kDa peak in Figure 3, black trace); lane 3, purified sample of the Ras-nanodisc conjugate (from the 200 kDa peak in Figure 3, red trace); lane 4, final NMR sample of Ras-nanodisc complex (appropriately diluted); lane 5, markers; lane 6, purified Ras used for conjugation (shown as a reference).

To demonstrate the native conformation of Ras at the surface of nanodiscs, we tested its nucleotide-binding function: performed nucleotide exchange of the endogenous GDP in the nanodisc sample for the fluorescent GDP derivative. The nucleotide exchange was induced in Ras-nanodisc conjugates with 1mM EDTA in the presence of 0.1 mM mant-GDP followed by gel-filtration to separate unbound mant-GDP from nanodiscs. Figure 5 shows fluorescence in the samples eluting from the gel-filtration column in the 200 kDa range when nanodiscs contained Ras (red) versus a control without a conjugated protein (blue). The characteristic emission peak at 440 nm indicates that Ras-nanodisc sample selectively retained mant-GDP while empty nanodiscs did not.

**Figure 5.**
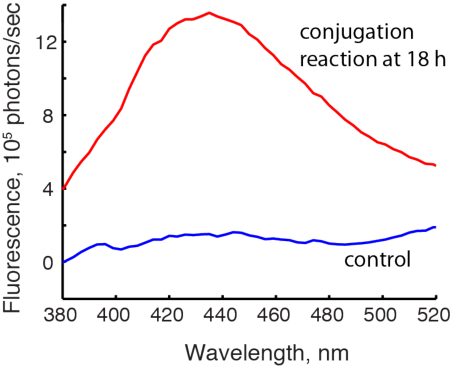
Demonstration of nucleotide-binding capacity of Ras-nanodisc conjugates. An aliquot from 200 kDa fraction shows mant-GDP fluorescence (red trace) while the control sample with unreactive nanodiscs does not (blue trace). Excitation wavelength: 360 nm.

## Results and Discussion

For NMR spectroscopy experiments with the ^15^N-labelled Ras-nanodisc, the fractions corresponding to the 200 kDa-peak were pooled and concentrated in pH 7.2 buffer including 170 mM NaCl (similar to the physiological ionic strength). We noted that nanodisc sample did not tolerate reduction of the ionic strength leading to precipitation in the concentrator if diluted with a low-salt buffer. Therefore, a moderately high salt content was chosen to ensure the sample stability.

Figure 6 demonstrates the ^15^N-^1^H 2D HSQC spectrum of the nanodisc-conjugated Ras (red) overlaid with the corresponding spectrum of GDP-bound Ras 1-166 (blue). Both spectra show similar chemical shift dispersion as well as overlap of many characteristic peaks (for example, strongly downfield-shifted resonances of G13 and K16). Spectral similarity indicates that Ras conjugated to a nanodisc is folded into a conformation similar to the structure of the G domain bound to GDP. Recording peak intensities as a function of time at the experimental temperature revealed that the sample is sufficiently stable. For example, spectral intensity remained nearly constant over the course of 12 hours at 30°C for a number of peaks (see Supporting Figure 1).

**Figure 6.**
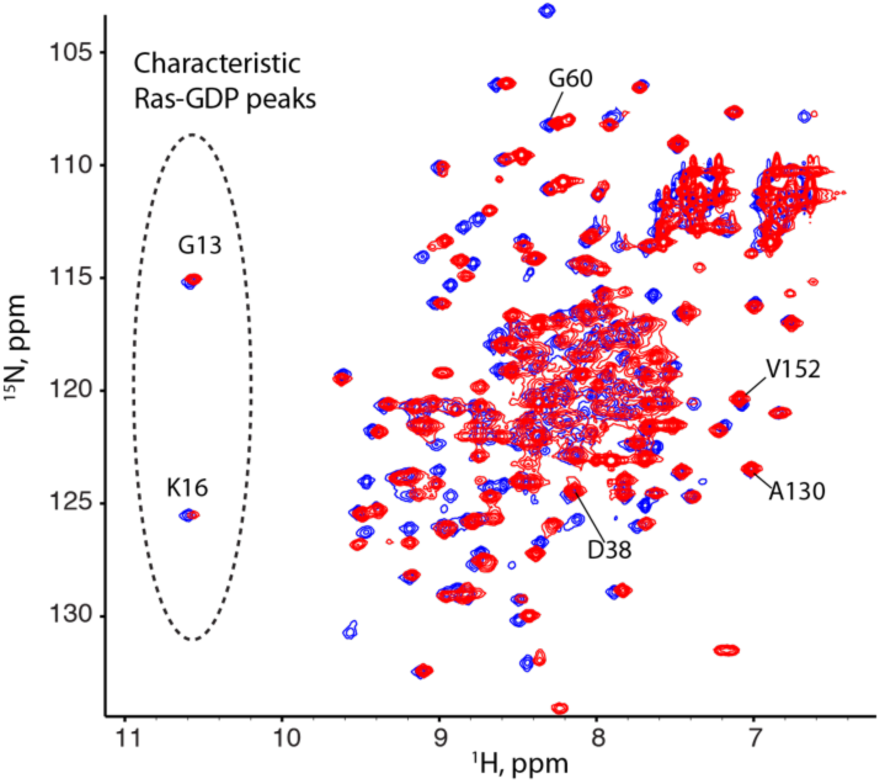
^15^N-^1^H HSQC of 0.14 mM Ras(1-181)-GDP-nanodisc sample (red) obtained at 30°C on 600 MHz Varian VNMRS instrument equipped with the Cold Probe. The spectrum of a truncated G domain, H-Ras(1-166)-GDP, is shown in blue for reference. Selected peaks are labeled and their time course is shown in Supporting Figure 1.

Integrity of the samples was also evaluated over the entire time of the NMR experiments using mass spectroscopy. Supporting Figure 2 shows MALDI-TOF analysis of the samples taken before and after all NMR experiments followed by a month-long storage at 4°C (2 months of total time) revealing minimal sample degradation.

Analysis of peak intensities before and after NMR experiments with Ras-nanodisc samples at different temperatures allowed to determine a relative rate of signal loss (due to precipitation) during NMR acquisition. Table 1 shows analysis of the sample life versus signal-to-noise ratio, S/N, at the corresponding temperatures. The sample precipitation rate was observed to increase approximately proportionally to the increase of S/N with temperature. Considering the cost of NMR time and other pathways of sample loss (proteolysis), the 37°C appears to be most appropriate condition (being also the physiologically relevant one for the human protein).

**Table 1.** Expected sample life vs. acquisition time to achieve the same signal-to-noise ratio at different temperatures.

| Temperature, -C | 20 | 30 | 37 |
| --- | --- | --- | --- |
| Rate of signal intensity decrease, %/hour | 0.07 | 0.33 | 0.6 |
| Projected sample life (using linear slope) | 60 days | 12 days | 6 days |
| Normalized sample life: | 10 | 2 | 1 |
| Normalized Acquisition time to achieve the same S/N at all temperatures | 11 | 3.7 | 1 |

One characteristic of the Ras-nanodisc complexes that is important to consider is *homogeneity* of the Ras-nanodisc ratio in the sample. Here we used 1:1 molar ratio of protein to lipid nanodiscs, which is 1:2 ratio of the protein to the nanodisc faces. If a high ratio of Ras to nanodisc is used to accelerate the reaction and increase the yield of the conjugate, it is likely to create some fraction of nanodiscs with two Ras molecules attached *to the same face* of the nanodisc, which will complicate measurement results. For example, Mazhab-Jafari and coworkers^*21*^ used 1:1 molar ratio of Rheb *to nanodiscs faces*, which is 2:1 ratio of protein to nanodiscs, and measured distances from protein nuclear spins to the lipid surface (using paramagnetic relaxation enhancement induced by a Gd^3+^-conjugated lipid). They found two sets of distances that were not consistent with a single orientation of Rheb at the nanodisc. The observed inconsistency led authors to propose that G domain of Rheb contacts the lipid bilayer in two distinct orientations. However, this result might also have been a manifestation of two populations of nanodiscs in the sample: one subpopulation with a single protein molecule per nanodisc face, while another (smaller) subpopulation—with two proteins at the same face, forced to assume different orientations simply due to spatial restriction.

To avoid ambiguities due to over-population of nanodiscs with conjugated Ras molecules, it is advisable to use significant molar excess of nanodisc faces over Ras as in our study (2:1) or greater. This approach guarantees that most nanodiscs carry only one protein molecule per face, yet creates samples with many “extra” empty nanodiscs. The empty nanodiscs may be removed if Ras and MSP protein carry orthogonal affinity tags.

## Conclusions

In this paper, we reported preparation and analysis of the uniformly ^15^N-labeled NMR samples of H-Ras conjugated to lipid nanodiscs. We found that Ras-nanodisc conjugation is a relatively reliable procedure resulting in moderately stable samples suitable for NMR spectroscopy. We stressed that achieving homogeneous ratio of protein to nanodiscs is essential for unambiguous data analysis.

## Acknowledgement

ELK acknowledges the Regular Research Grant 2012 from Committee on Research (COR), Marquette University.

## Supporting Information

**Supporting Figure 1.**
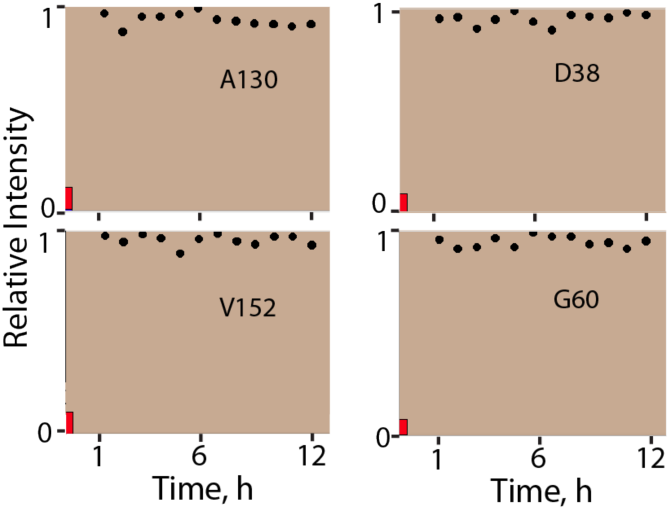
Assessment of stability of the protein-nanodisc complex. Normalized peak intensity for a few randomly chosen peaks (labeled in A for visual aid) plotted versus time at 30°C. Graphs were created in Sparky using “es” extension from *Extension:Kovrigin* custom sub-menu. Vertical red bar indicates 3x noise RMSD for a relative scale. Custom Sparky extension code is available upon request.

**Supporting Figure 2.**
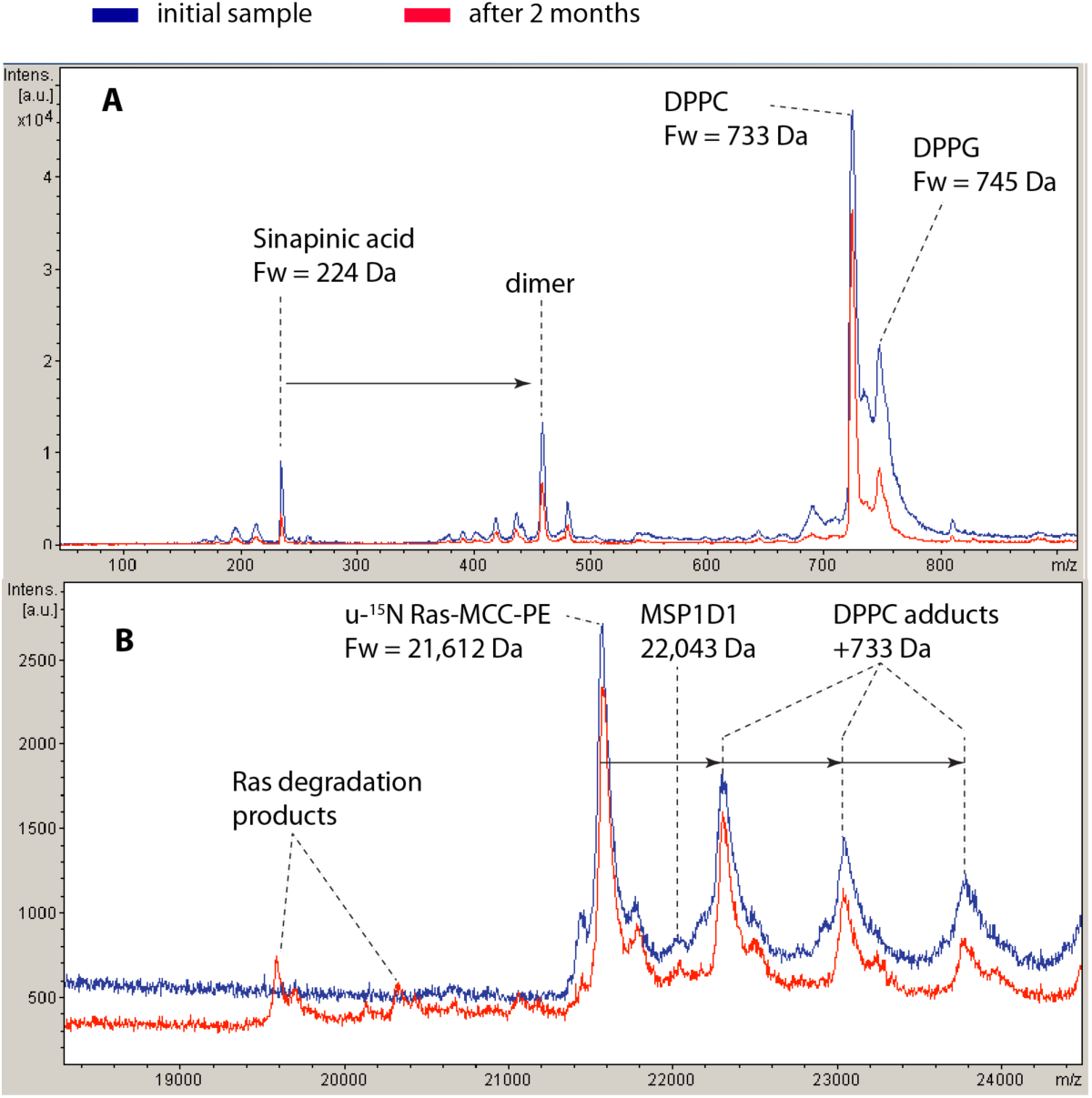
MALDI-TOF mass spectra of a Ras-nanodisc sample. (A) Low-m/z range; (B) high m/z range. Blue trace, Ras-nanodisc sample prior to NMR measurements; red trace, the same sample after 2 months at variable temperatures: 200 hours at 37°C, about 20 days at 20°C, the rest at 4°C. Calculated formula weights expected for Ras conjugate and other molecules are given in the Figure. MSP1D1 protein did not ionize well and appeared with a relatively low intensity.

